# Climate change risk to southern African wild food plants

**DOI:** 10.1101/2020.08.22.262535

**Authors:** Carina Wessels, Cory Merow, Christopher H. Trisos

## Abstract

Climate change is a threat to food security. Wild-harvested food plants (WFPs) are important for the diets of millions of people and contribute to food security, especially in rural and low-income communities, but little is known about climate change risk to these species. Using species distribution models, we examined climate change risk to 1190 WFP species used by 19 native language groups in southern Africa. We project that 40% of species will experience a decrease in range extent within southern Africa by 2060–2080 under a low warming scenario (Representative Concentration Pathway (RCP) 2.6), increasing to 66% under a high warming scenario (RCP 8.5). Decreases in geographic range are projected for >70% of WFP species traditionally used by some language groups. Loss of suitable climatic conditions is projected to decrease WFP species richness most in north-eastern southern Africa – with losses of >200 species – while increases in species richness are projected in the south and east of South Africa. Availability of WFP species for food security during lean times is also projected to change. Specifically, in south-eastern South Africa, local diversity of WFPs is projected to increase, while maize and sorghum yields decrease. However, this potential WFP nutritional safety net may be lost in central parts of the region, where declines in both crop yield and WFPs are projected. By looking beyond conventional crops to the exceptional diversity of WFPs, this research makes a first step towards understanding the linkages between WFPs, traditional knowledge, agriculture, food security, and climate change.

## Introduction

With the world population soon to reach eight billion people, achieving global food security is a complex challenge. This challenge is exacerbated by climate change (Hoegh-Guldberg et al. 2018). Research describing climate impacts in terms of food security has mostly focused on the effect of climate change on agriculture, mainly on a small number of staple crops (Vermeuelen et al. 2012; Challinor et al. 2014; Rosenzweig et al. 2014; Wu et al. 2014; Springmann et al. 2016). Changes in temperature and rainfall have already negatively impacted crop yields (Carleton and Hsiang, 2016; Ketiem et al. 2017) and future projections indicate substantial reductions in yields and nutritional quality of cereal crops globally for 2°C of global warming, with particularly severe impacts on maize in sub-Saharan Africa (Vermeulen et al. 2012; Porter et al. 2014, Hoegh-Guldberg et al. 2018). Apart from agriculture, food security research has further concentrated on fisheries, despite the fact that other wild-caught or wild-harvested foods also play a vital role in food security, especially in Africa and other developing regions (Bennet and Robinson 2000; Bharucha and Pretty 2010; Hickey et al. 2016; Rowland et al. 2017). Wild foods, including both plant and animal products, contribute to food security via direct consumption either on a regular basis or in times of shortage, and when sold for income that could be reinvested in other food purchases (FAO 2019). Wild foods can thus be a potential solution to help overcome food insecurity (Shaheen et al. 2017).

Wild food plants (WFPs) in particular, provide pronounced dietary diversity and can make an important contribution to micronutrient intake (Grivetti and Ogle 2000). For many people in sub-Saharan Africa, WFPs are already an essential livelihood safety net when other sources of food fail, such as during times of drought or other climate impacts (Vinceti et al. 2013; Wunder et al. 2014; Shumsky et al. 2015). For example, in the arid Ferlo region of Senegal, the specific wild species consumed during seasonal lean periods are crucial contributors of vitamins A, B2 and C (Becker 1983) and in the Sahel, numerous desert WFPs are sources of essential fatty acids, calcium, iron and zinc (Glew et al. 1997). WFPs consumed during the lean dry season in Nigeria were also found to have a higher micronutrient and energy content, compared to WFPs used in the productive wet season (Lockett et al. 2000). In lean times in Zimbabwe, the quantities of wild fruits consumed and sold to generate income for additional food expenses also increases in poor rural households (Mithöfer and Waibel 2004). In Botswana, when drought leads to crop failure, rural communities rely on WFPs to sustain their livelihoods until conditions improve (Twyman 2001; Ohiokpehai 2003; Mojeremane and Tshwenyane 2004). In Mali, Tanzania, and Zambia, household surveys of climate change adaptation strategies showed that forest products (which includes wild foods) can play a crucial role in reducing household vulnerability by providing alternative sources of food and income during droughts and floods (Robledo et al. 2012). However, this adaptation response may be at risk. For instance, in Benin, the economic value of three important wild tree species – *Adansonia digitata* (baobab), *Parkia biglobosa* (African locust bean tree), and *Vitellaria paradoxa* (shea tree) – is projected to decrease up to 50% by 2050 under a high warming scenario (Heubes et al. 2012). Assessing WFPs as a climate change adaptation strategy could also be an important example of the potential to bridge western scientific and traditional knowledge systems. The use of traditional knowledge for climate change adaptation is potentially a key strength for African communities (Makondo and Thomas 2018).

Despite these indications that WFPs could increase resilience to climate change, limited research has been done on climate change risks to wild-harvested food plants. Given the projected impact of global climate change on food security, assessing climate change risk to WFPs is necessary for climate change adaptation planning, especially in the most vulnerable regions of the world. Thus, the primary aim of this study was to determine where in southern Africa WFPs occur and whether these WFPs will be threatened by climate change. A secondary aim was to determine if regions where WFPs are projected to be at risk in the future overlap with regions of projected agricultural crop yield loss under climate change, in order to assess whether WFPs may be available as a coping strategy for communities facing negative impacts on crop yields. Knowing the projected impact of climate change on WFPs could afford valuable time for people to adapt to changing circumstances, especially those communities that already rely on WFPs, or are vulnerable due to crop yield losses, or both. This research could also aid in developing advance warning systems that are representative of a larger range of food supply options in a region in order to mitigate risk through targeted food security and species conservation management actions.

## Materials and Methods

### Wild food plant species

To identify WFPs we used a recently published list of 1740 southern African edible plant species (Welcome and Van Wyk 2019). The list includes food plants from Botswana, Eswatini (formerly Swaziland), Lesotho, Namibia, and South Africa. It is the most complete inventory of wild edible plants in southern Africa to date, and one of the most comprehensive globally for known usage of WFPs by local communities. We focused on WFP species native to southern Africa. Exotic species make up 15% of the edible plant species listed for the region (e.g. naturalised weeds used as vegetables), but we excluded these species due to limits in modelling the global range extent of exotic species. All native WFP species names (n=1479) were run through The Taxonomic Name Resolution Service (http://tnrs.iplantcollaborative.org) to match with species names for species occurrence record data, by checking for alternative spellings and converting synonyms to current accepted names. We also obtained information from the inventory on how each plant is utilised, and when adequate information was available, which indigenous language/ethnic groups make use of WFPs in southern Africa (see Supplementary Table 1 for a list of uses and language groups). WFP uses can be particular to different language/ethnic groups and are often closely connected with traditional knowledge in these groups (Welcome and Van Wyk 2019, 2020).

### Species occurrence records

Wild food plant occurrence records were obtained from the Botanical Information and Ecology Network (BIEN) (http://bien.nceas.ucsb.edu/bien/). Although the WFP data is restricted to southern Africa, we considered the full extent of the geographic range of occurrences for each species in order to more accurately reflect the species environmental niches and projected response to future climate change. Occurrence records were passed through BIEN’s geovalidation filters to remove records with errant coordinates and check that the latitude and longitude of a recorded observation falls within its specific declared political divisions (http://bien.nceas.ucsb.edu/bien/biendata/bien-3/validation/). A number of steps were performed to prepare occurrence records before modelling. Firstly, for species with more than 20 occurrences, occurrences were thinned to ensure that all retained records were at least 20 km from one another in order to reduce spatial autocorrelation (Aiello-Lammens et al. 2015). Secondly, outliers in geographic and environmental space were discarded based on a Grubb’s outlier test with p=0.0001 (Grubbs 1950) (implemented with the *R* package outliers (Komsta 2011)). Finally, presences were clustered into five folds spatially stratified for cross-validation. Only species with 10 or more records were retained and used in subsequent analyses (n=1190, representing 80% of identified native WFPs in the southern African region).

### Historical climate and environmental data

Environmental predictors included six climate and five soil layers. 30-year time-averages of historical climate data were downloaded from WorldClim version 2.0, at a spatial resolution of 10 km, for the period 1970–2000 (Fick and Hijmans 2017). To reduce collinearity among climate variables in the species distribution models, four bioclim variables were chosen from the full set of 19 bioclim variables, based on their low correlation with each other (r < 0.7): mean annual temperature, mean diurnal temperature range, annual precipitation, and precipitation seasonality. Two additional variables were added based on expert recommendation and since they also had a correlation coefficient of r < 0.7 with the four selected bioclim predictors. These were (1) aridity, calculated as the maximum accumulated water deficit and (2) precipitation in the warmest quarter divided by the sum of precipitation in the warmest quarter and precipitation in the coldest quarter. Five soil layers were obtained from https://soilgrids.org. These were bedrock depth, mean bulk density, mean pH, proportion silt, and proportion clay in the first four soil horizons. To generate the soil layers, the 250 m resolution layers available on soilgrids.org were aggregated to the 10 km grid defining the climate layers. These aggregated layers correlated at r < 0.7 with the climate layers.

The set of 11 predictor variables was the starting point for modelling the distribution of each species. Within each species-specific model domain, however, the 11 predictors were further subset to ensure they had r < 0.7 within the modelling domain by retaining the largest subset of predictors below this correlation level. This emphasis on removing correlated predictors for each species was done to reduce the influence of correlations among predictor variables on the fitted models, when projecting the future range of each species.

### Future climate scenarios

Future climate data for 20-year time averages were downloaded from WorldClim version 1.4 (https://worldclim.org/data/v1.4/cmip5.html), at a spatial resolution of 10 km, for the period 2061–2080. Future climate projections from seven Coupled Model Intercomparison Project (CMIP5) Global Climate Models (GCMs), that were downscaled and calibrated by WorldClim (Hijmans et al. 2005), were used in this study (Supplementary Table 2). Data were downloaded for two future greenhouse gas scenarios: representative concentration pathway (RCP) 2.6 and RCP 8.5 (van Vuuren et al. 2011). RCP 2.6 is a low emissions scenario, likely keeping global warming below 2 °C by 2081–2100 compared to 1850–1900, whereas RCP 8.5 has been characterised as an extreme high emissions scenario representing more than 4 °C of global warming by 2081-2100 (Cubasch et al. 2013).

### Species distribution modelling

Inhomogeneous Poisson Point Process (PPM) models (Warton and Shepherd 2010; Renner et al. 2015) were used to obtain the historical range of species and subsequently to make spatial projections for future climate scenarios. Future distributions, under RCP 2.6 and RCP 8.5, were predicted for every species for each of the seven different GCMs.

PPMs are a generalisation of the MaxEnt algorithm, which is commonly used where only presence data are available (Phillips et al. 2006). PPM settings were chosen to balance underfitting (that is, excessively smooth models that over predict range size) with overfitting that would underestimate species range sizes. PPM models were fitted using regularized down-weighted Poisson regression (Renner et al. 2015) in the *R* package *glmnet* (Friedman et al. 2010). Different feature classes of increasing complexity were added depending on the number of occurrence records available for each species: linear and quadratic for all species, and product for species with more than 100 records. When models failed to converge with more complex feature classes the next simpler feature set in this hierarchy was used. Linear and quadratic features were always paired together to ensure modal responses were always included as an option, in order to reflect that species often exhibit modal responses to environmental gradients (Austin 2007). In cases where linear and quadratic features failed, the number of predictor variables was reduced by selecting the five (or three, if five failed) predictors with the highest univariate correlations with presence data, in order to generate linear and quadratic features. This sequential approach of applying simpler features increases the robustness of the modelling approach to idiosyncrasies in data for individual species. This is in order to ensure that models could be fit for each species with 10 or more presence records.

The regularisation parameter was determined based on 5-fold cross-validation with each fold, choosing a value one standard deviation below the minimum deviance (Hastie et al. 2009). The resulting five models for each species were combined in an unweighted ensemble, which is interpreted as a relative occurrence rate (ROR) (Fithian and Hastie 2013; Merow et al. 2013). Continuous predictions of ROR were converted to binary absence/presence predictions for each species by choosing a threshold based on the fifth percentile of training presence locations.

When projecting species occurrences using future climate scenarios, extrapolation in the species distribution model was limited to one standard deviation beyond the data range of the observed occurrence records for each predictor variable. This approach was used to allow for a small amount of extrapolation beyond the modelled limits of the observed species niche, while also limiting the influence of monotonically increasing marginal responses. The latter can lead to statistically unsupported and likely biologically unrealistic projections of species responses to climate change.

Due to lack of data on dispersal distances, species-specific dispersal capacity was not included in modelling changes in species geographic range in response to future climate change scenarios. Instead, spatial predictions of future occurrences were constrained to be in the WWF ecoregions (Dinerstein et al. 2017) where a species occurs currently, and the immediate neighbours of those ecoregions. Although ecoregions differ in size, this approach prevents predictions of future occurrences being extreme distances away from the current range of each species. It is preferred to a strict spatial rule because it integrates the high degree of biome-level niche conservatism found in southern African plant species ranges (Crisp et al. 2009) into the modelled dispersal constraint.

### Analysis of model outputs

For every species, we made majority consensus maps for future species distributions under RCP 2.6 and RCP 8.5. This was done by only assigning a species as present if a majority (four out of seven) of the species distribution models from the GCMs agreed on the species being present in a given 10 km grid cell. We calculated the projected change in species richness for each grid cell under each scenario, by adding up the number of species presences in each grid cell for the historical time period (1970–2000) and each future scenario, and then subtracting the historical value from the future value.

For each species, we also calculated the percentage change in geographic range extent between future and historical modelled geographic ranges. We calculated the mean geographic range change for each species across the seven GCMs, and additionally calculated the range change by category of plant use, as well as by indigenous language group use.

### Crop yields and wild food plants

In order to determine whether the projected change in WFP species richness overlaps with regions where agricultural crop yield losses are projected due to climate change, we downloaded crop yield projections for maize (*Zea mays*) and sorghum (*Sorghum bicolor*). Data were downloaded from The Inter-Sectoral Impact Model Intercomparison Project (ISIMIP) (www.isimip.org) under the ISIMIP Fast Track simulation protocol, as this was the only protocol with RCP 8.5 data available. We chose maize and sorghum because these are two of the most important agricultural crops for small scale farmers in southern Africa (Zinyengere et al. 2014). Maize yield data were downloaded for four crop models, and sorghum yield data for two crop models (Supplementary Table 3). Each crop model had projections for five different GCMs (Supplementary Table 3), totalling 20 model runs for maize and 10 model runs for sorghum, for each of the historical conditions, RCP 2.6, and RCP 8.5. Crop yield rasters were downloaded at 0.5°x0.5° regular global grid resolution, with CO_2_ fertilisation, no irrigation and Shared Socioeconomic Pathway (SSP) scenario two. SSP2 is a “middle of the road” scenario of projected socioeconomic global change up to 2100, where trends are expected to follow their historical patterns, with medium challenges to adaptation and mitigation (Riahi et al. 2017). We chose a no irrigation scenario because most small-holder farmers in southern Africa depend only on rain for crop production and this would likely include those communities that make use of WFPs in times of hardship. We retained only annual crop yield projections in years that matched historical (1970–2000) and future (2061–2080) climate time periods available for WFPs. For each crop model, we first calculated the mean of the yield data for the five available GCMs, and then the mean across crop models to get the overall mean crop yield for maize and sorghum for historical conditions, RCP 2.6, and RCP 8.5. To enable comparison of projected changes in crop yield and WFP species richness, WFP projections were regridded using nearest neighbour interpolation to match the lower 0.5° spatial resolution of the crop yield data. We mapped the percentage change between future and historical crop yield and used these maps, together with maps of percentage change in WFP species richness, to identify regions where both crops and WFPs, or just one of these are at risk. For this analysis, only WFPs that are used as a snack, fresh or in situ (n=686), cooked vegetables (n=407), ground as flour (n=91), or as famine food (n=64; famine food is a term used for a category of food plant that is not really eaten except under conditions of famine, to avoid starvation) were included (Welcome and Van Wyk 2019). This selection of WFPs represents 81% of the total number of species used in this study. Wild food plants in these categories of use were chosen as they are the most likely to replace staple crops, such as maize and sorghum, when these crops fail.

## Results

### Risk to wild food plant species

WFP species richness in southern Africa generally increases from west to east across the region with the Eastern Cape, Kwazulu-Natal and Mpumalanga provinces of South Africa having the highest species richness (Figure 1A). Within the western part of the region, there is also a north-south gradient of increasing WFP species richness from more arid habitats in Namibia to the Cape Floristic Region in the Mediterranean climate zone of South Africa. These spatial patterns in WFP species richness stay the same when looking at individual categories of the uses of particular WFPs most important for food security and as nutritional supplements (Figure 1B–E).

**Figure 1.**
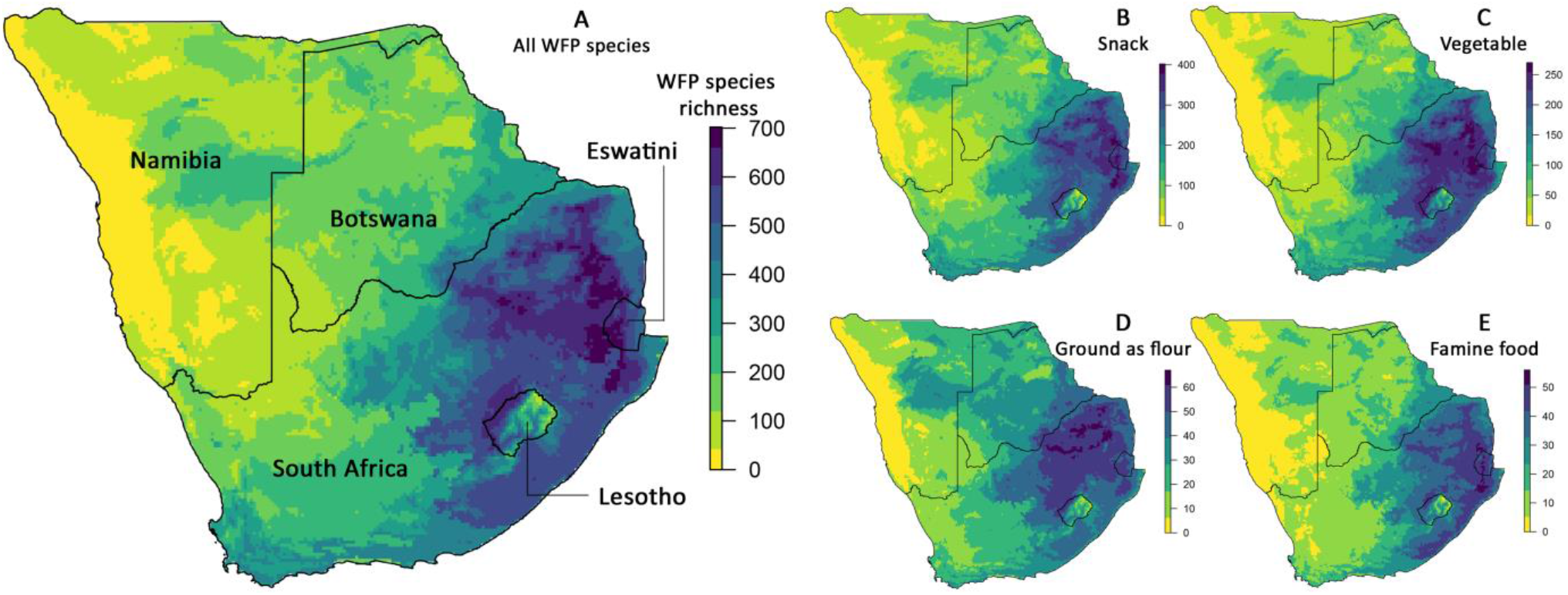
Wild food plant (WFP) species richness in five southern African countries: Botswana, Eswatini, Lesotho, Namibia, and South Africa. A – all WFP species modelled in this study (n=1190); B – species eaten as a snack, fresh or in situ (n=686); C – species eaten as cooked vegetables (n=407); D – species that are ground as flour (n=91); E – species used as famine food in times of hardship (n=64).

We project that under RCP 2.6, a low emissions scenario, 40% of the 1190 native WFP species modelled will experience a decrease in geographic range extent by 2060–2080 within southern Africa (Figure 2A): 38% are expected to lose up to a quarter of their range, 2% lose between a quarter and half their range, and less than 1% lose more than half of their range (mean range loss = 9% ± 10 SD; Figure 2A). However, species gaining range extent within southern Africa outnumber those projected to have range losses, with a total of 60% expected to experience a range increase under RCP 2.6 (Figure 2A): 40% are expected to gain up to 25% in geographic range area, while 20% of species gain more than 25% in range (Figure 2A). No regional species extinctions, estimated as more than 90% range loss, were projected under RCP 2.6.

**Figure 2.**
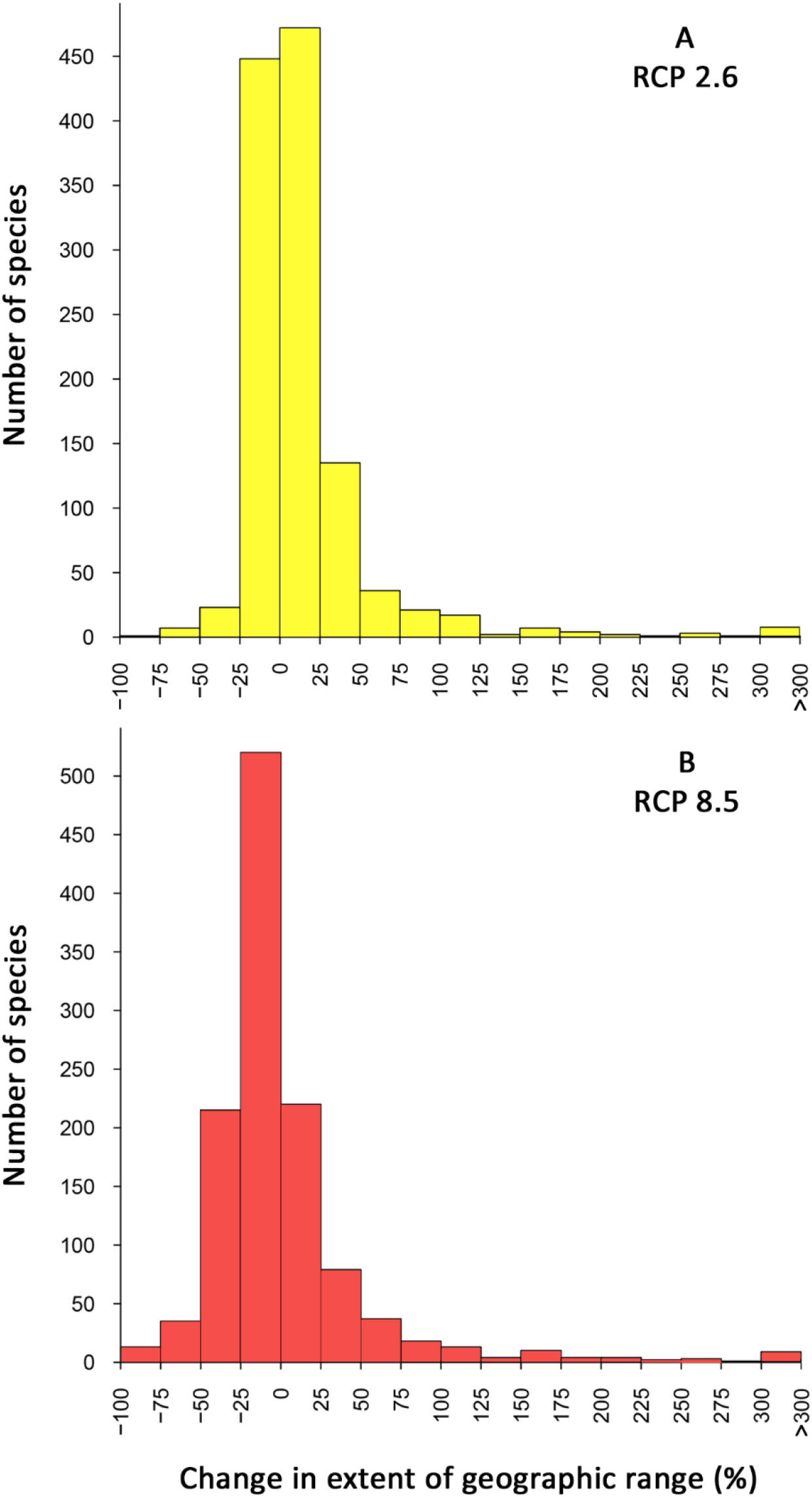
The mean change in extent of geographic range (%) for 1190 southern African native wild food plant species calculated from seven Global Climate Models under: (A) a low warming scenario (RCP 2.6), and (B) a high warming scenario (RCP 8.5).

In contrast, under RCP 8.5, a high emissions scenario, the pattern of projected range change is reversed, with a majority (66%) of native WFP species projected to suffer a decrease in range extent within southern Africa (Figure 2B): 44% are expected to lose up to a quarter of their range, while 18% lose between a quarter and half, and 4% of species lose more than half their range extent (mean range loss = 22% ± 17 SD; Figure 2B). Only 34% of species are projected to experience a range increase under RCP 8.5: 19% are expected to gain up to 25% in range area, and 15% gain more than 25% in range (Figure 2B). Five regional species extinctions, estimated as more than 90% range loss, were projected under RCP 8.5. These species are *Combretum engleri*, used as a snack; *Euphorbia inermis*, used as a snack and cooked vegetable; *Grewia schinzii,* used as a snack and non-alcoholic beverage; *Searsia horrida*, used as a snack and to curdle milk; *Vachellia hebeclada*, used as a snack.

Projections of changes in species richness under RCP 2.6 show that the loss of suitable climatic conditions is expected to decrease species richness of WFPs most in the north-eastern parts of southern Africa with local extinctions of more than 100 species (Figures 3A and B). This pattern is intensified under RCP 8.5 with multiple regions in north-eastern South Africa projected to have decreases of more than 200 species and local species losses of more than 50% of WFPs are projected for most of Botswana (Figures 3C and D). In contrast, increases in local species richness are projected in more southern and eastern regions due to species shifting their ranges southward and into cooler montane regions under both RCP 2.6 and RCP 8.5 (Figures 3B and D). The largest relative local species richness increases are projected in the western parts of southern Africa, with increases of more than 270% on the west coast of Namibia, although this represents a small change in absolute species numbers – 12 species under RCP 2.6 and 14 species under RCP 8.5 (Figures 3A and C). Large increases in local species richness of between 25–125% are also projected in the south and south-eastern regions of South Africa (Figures 3A and C).

**Figure 3.**
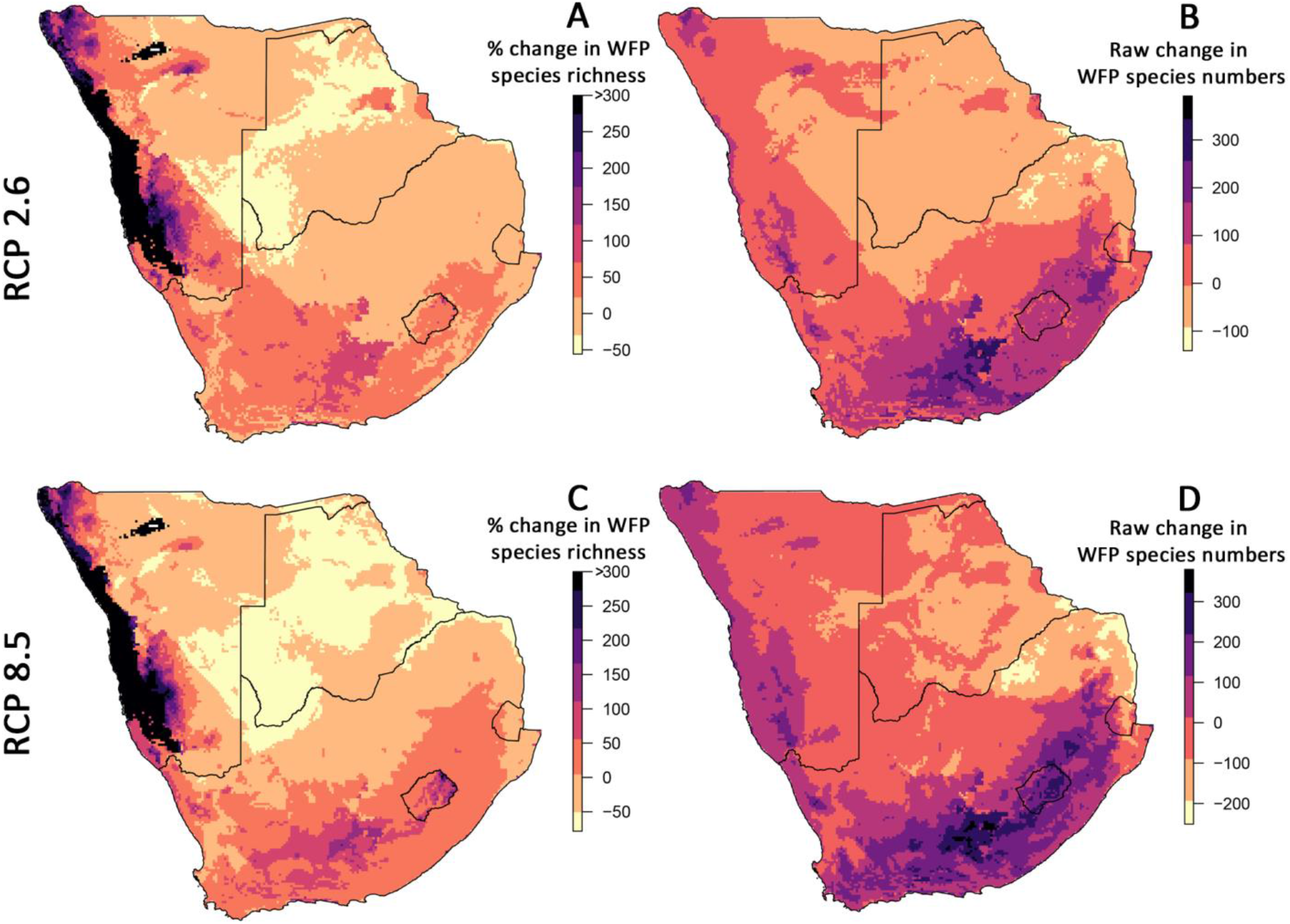
The projected change in wild food plant (WFP) species richness for a low warming (RCP 2.6) and a high warming (RCP 8.5) scenario. A and C show projected percentage change in species richness, whereas B and D show the raw change in WFP species numbers.

### Risk by wild food plant usage and language group

All categories of WFP use – except for species used as syrup, to curdle milk, or as yeast – are projected to have more species undergo increases than decreases in range extent under RCP 2.6 (Figure 4; Supplementary Table 4). In contrast, under RCP 8.5 decreases in range area are projected for over half of species in every use category except alcoholic beverages (45% of species), with some categories projected to be more negatively affected than others (Figure 4; Supplementary Table 4). For example, substantial range decreases are projected for species used as milk curdlers (74%), sources of moisture/thirst quenchers (77%), preservatives (73%), thirst or appetite suppressants (91%), syrup (83%), tea substitutes (80%), and as yeast (78%) (Supplementary Table 4). When looking at the categories of use of WFPs most important for food security and as nutritional supplements, 59–69% of species are projected to experience decreases in range under RCP 8.5, including species used as: cooked vegetables (69%), as famine food (59%), ground as flour (67%), and eaten as snacks, fresh or in situ (63%) (Supplementary Table 4).

**Figure 4.**
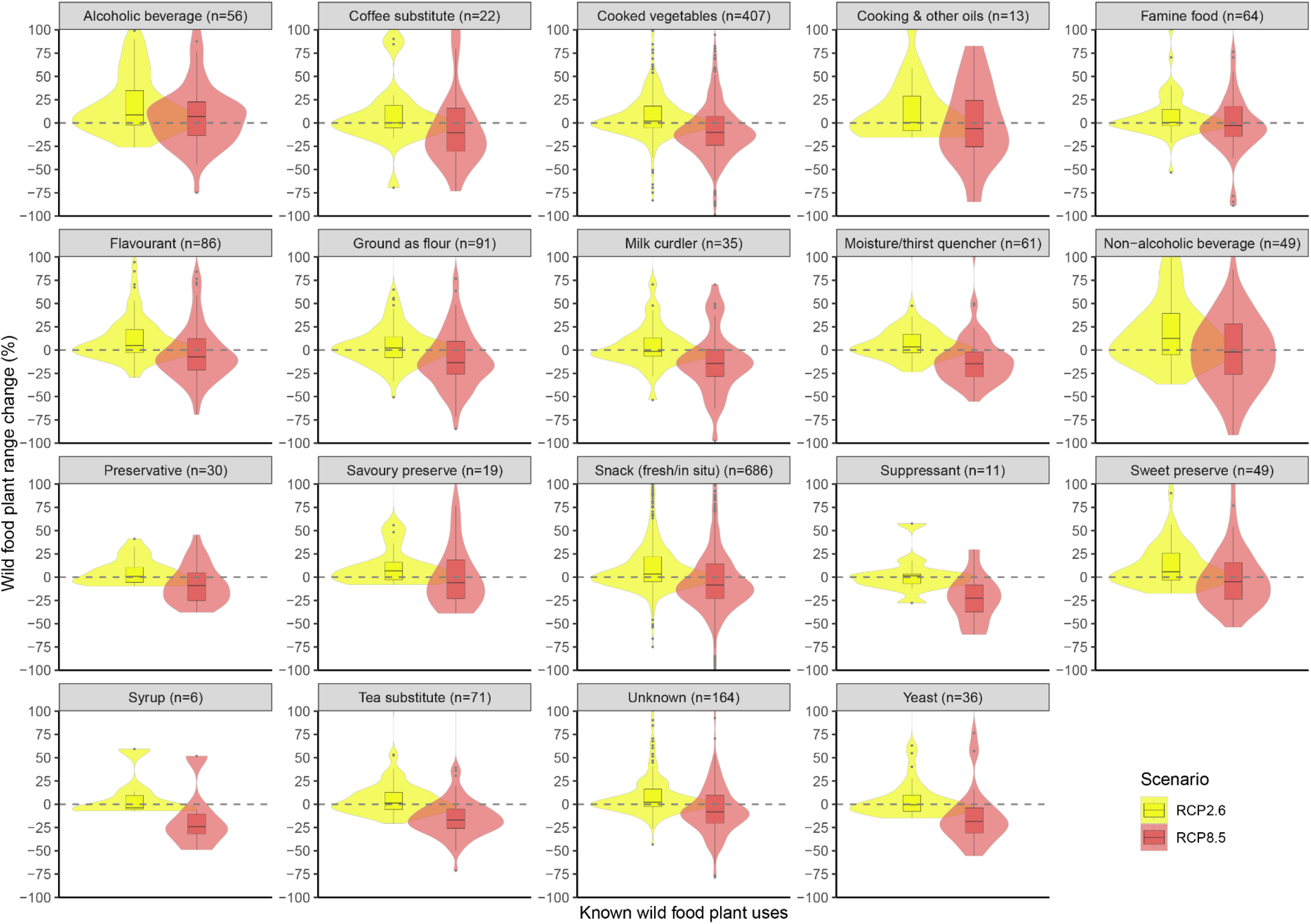
Projected percentage change in wild food plant range extent by category of plant use for RCP 2.6 (yellow) and RCP 8.5 (red). Only range increases of up to 100% are shown, as most of the species range change lies within this limit (see figure 2). The median, 25^th^ and 75^th^ percentiles are shown by box plots within the violin plots. Potential outliers are indicated by grey dots. The numbers of species (n) are shown for each category of use.

WFPs are an important component of the indigenous and local knowledge of food systems, and their use in southern Africa varies by language/ethnic group (Welcome and Van Wyk 2019; 2020). For RCP 2.6, over half of species used by each language group are projected to experience range increases, except for species used by the Xóõ language group, from the dry Kalahari region of Namibia, Botswana, and South Africa (49% of species) (Figure 5; Supplementary Table 5). However, under RCP 8.5, range decreases are projected for more than 50% of species used by every language group, except the Lozi from the Caprivi Strip in Namibia and northern Botswana (47% of species) (Figure 5; Supplementary Table 5). Some language groups are expected to be especially negatively affected, specifically, geographic range decreases are projected for over 76% of species used by the Southern Sotho, 71% of species used by the Xhosa, and 74% of species used by the Xóõ.

**Figure 5.**
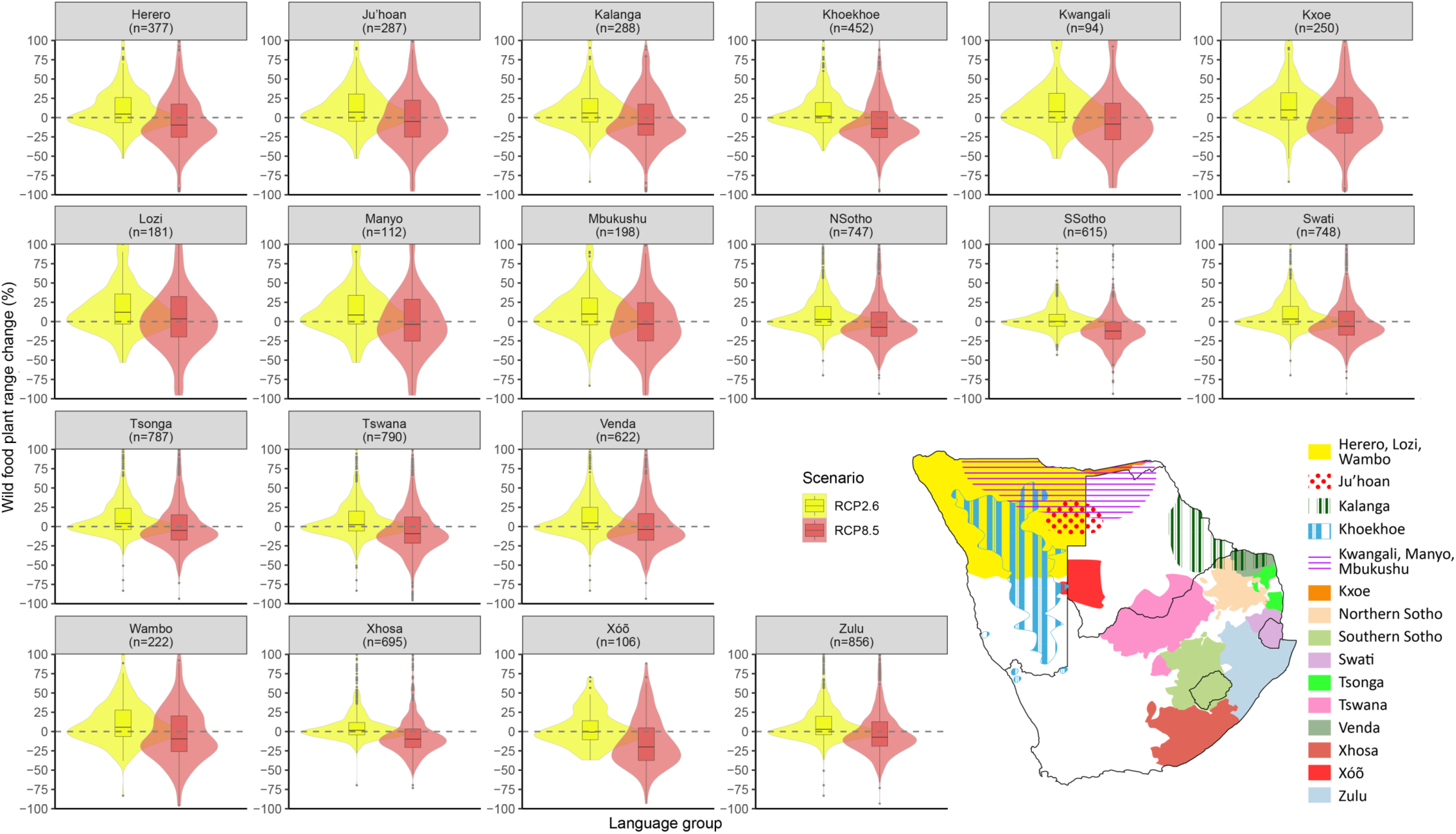
Projected percentage change in wild food plant range extent by category of known language group use for RCP 2.6 (yellow) and RCP 8.5 (red). Only range increases of up to 100% are shown, as most of the species range change lies within this limit (see figure 2). The median, 25^th^ and 75^th^ percentiles are shown by box plots within the violin plots. Potential outliers are indicated by grey dots. The numbers of species (n) are shown under each category of use. The map inset gives an indication of the distribution of the 19 language groups in southern Africa (data from Van Wyk et al. 2011).

### Intersecting risk to wild food plants and crops

People living in rural areas already use WFPs to supplement their diets, especially during times of hardship when staple crops such as maize and sorghum fail and WFPs might thus form part of climate change adaptation strategies for food security. To identify regions where both crop yields and WFPs, or just one of these are projected to be at risk from climate change, we compared climate change risks for maize and sorghum yields with risks for a selection of WFPs that are most likely to replace staple crops when these crops fail. This selection constituted 81% (n = 964) of the total number of WFPs. Our approach identified large regions of southern Africa where both crop yields and WFPs are at risk under future climate change, highlighting concerns for food security under both RCP 2.6 and RCP 8.5 (Figure 6). South Africa’s North West and Free State provinces, as well as parts of northern Namibia are projected to experience declines in both crop yield and WFP species richness. Under RCP 8.5, the highest risk to maize, sorghum, and WFPs is projected for the north-eastern border between South Africa and Botswana, as well as for sorghum and WFPs in northern Namibia and Eswatini (Figures 6B and D). In contrast, a decrease in crop yield with an increase in WFPs is projected for maize and sorghum in parts of south-east South Africa (Eastern Cape province), as well as for sorghum in the north of Namibia (Figure 6). Under RCP 2.6, a small region on the northernmost border of South Africa and Botswana is projected to experience an increase in crop yield with a decrease in WFPs (Figures 6A and C). Regions where increases are projected for both crop yield and WFP species richness are the southern and eastern parts of southern Africa, including Lesotho and parts of the Western Cape, Eastern Cape, Kwazulu-Natal and Mpumalanga provinces of South Africa (Figure 6).

**Figure 6.**
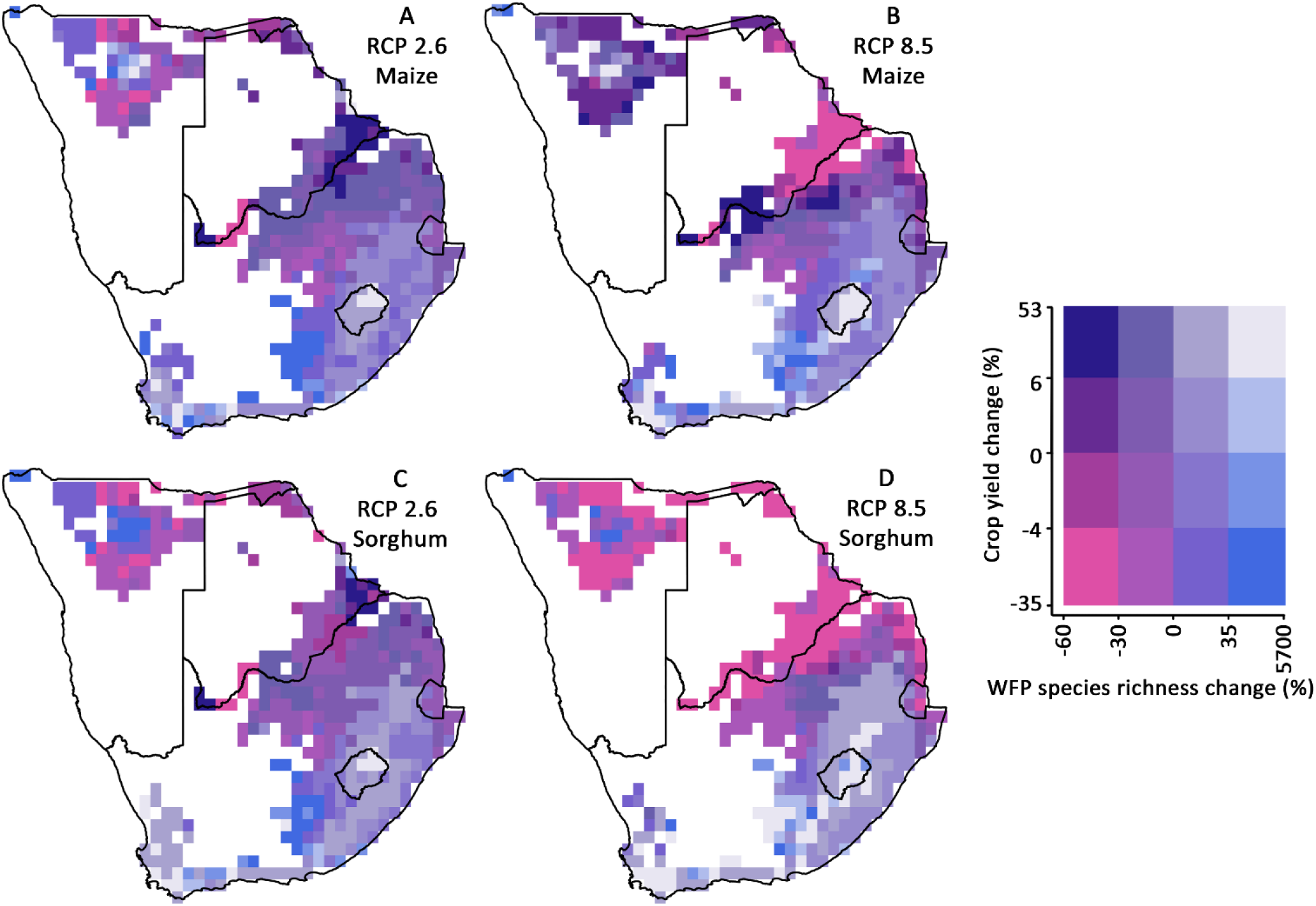
Projected crop yield change (%) combined with wild food plant (WFP) species richness change (%) for a low emissions (RCP 2.6) and high emissions (RCP 8.5) scenario in 2060–2080 compared to historical climate (1980–2000). Only WFPs that are used as snacks, fresh or in situ (n=686), cooked vegetables (n=407), ground as flour (n=91), or as famine food (n=64) were included in this analysis as they are the most likely to replace staple crops, such as maize and sorghum, when these crops fail. White regions are projected as unsuitable for maize and sorghum both in historical and future warming scenarios. Pink colours show regions where both crop yields and WFP species richness are projected to decrease.

## Discussion

Within southern Africa, large variations are projected in WFP species responses under both low and high warming scenarios, with range decreases expected for some species and increases for others. Under RCP 2.6, the less than 2 °C warming scenario, increases in range area within southern Africa are projected for 60% of WFP species and reductions for 40% of species. Range losses are projected to intensify for RCP 8.5, the greater than 4°C warming scenario, with 66% of species projected to experience range reductions and only 34% range increases (Figure 2). We note that because our assessment of changes in range area was restricted to southern Africa—the focus region for the WFP data—our projections do not reflect total change in range area for those species for which current ranges extend outside the region. This prevents the use of our projections for global extinction risk estimates, because widespread species projected to track suitable climate conditions southwards, may increase in range area within southern Africa, even as their overall range area in Africa decrease.

As the geographic ranges of WFP species change, the local diversity of WFPs, indicated by the number of species in a 10 km grid cell, is also projected to change. Under RCP 8.5, local species losses of more than 200 species are projected in the north-eastern parts of southern Africa, and reductions of more than 50% of current WFP species richness are projected for most of Botswana (Figure 3). Over these semi-arid regions of southern Africa, high warming rates and an increase in aridity are projected, with already water-stressed countries like Botswana and Namibia likely to get even hotter, drier and more water-stressed (Hoegh-Guldberg et al. 2018). In contrast, the largest relative increases in local WFP species richness are projected for the western parts of southern Africa, particularly the coastal region of Namibia, although this represents a small change in absolute numbers of species (Figure 3). Native species in the hot and arid west of southern Africa already have a high degree of stress tolerance (Midgley and Thuiller 2007) and it might be that as a result, WFP growth forms in the region, such as geophytes, have a higher resilience to climate change. These species are projected to persist under climate change, thereby maintaining current species numbers and leading to an increase in species richness under future climate conditions, as other species shift their ranges into more coastal regions of Namibia. A possible explanation for increased suitability of this region and increased WFP species richness in coastal Namibia, is the projected increase in precipitation through an upsurge in fog occurrence along the west coast (Haensler et al. 2011; Rohde et al. 2019).

Increases in WFP species richness are also projected in the southern and eastern regions of South Africa due to species shifting their ranges southward and into cooler montane regions under both RCP 2.6 and RCP 8.5 (Figure 3). In the east, wetter conditions are projected in the Drakensberg mountain range, a dominant feature in South Africa’s landscape, with an elevation of over 3000 m (Hewitson and Crane 2006; Engelbrecht et al. 2009). In the south, previous studies have projected that this region, which consists of mainly the Albany thicket, Grassland and shrubland semi-arid Nama Karoo biomes, will likely undergo substantial future changes in dominant plant growth forms in response to climate change, as well as due to increased atmospheric CO_2_ concentrations (Masubelele et al. 2015). For example, in response to a combination of increasing atmospheric CO_2_ and aridification, the Nama Karoo biome is projected to shift to the east into the Grassland biome (Ellery et al. 1991; Rutherford et al. 1996; Midgley et al. 2002; Midgley et al. 2008). For plants of drier climates, such as those that occur in the Nama Karoo, positive changes in water use efficiency, positive responses to increasing CO_2_ concentrations, and the potential for CO_2_ fertilisation, may be especially beneficial (Cowling 1999; Cowling and Sykes 1999; Thuiller et al. 2006).

WFP uses in southern Africa are often particular to different ethnic/language groups, and also closely connected with traditional knowledge (Welcome and Van Wyk 2019; 2020). Of the 19 southern African language groups examined by Welcome and Van Wyk (2019; 2020), the Zulu language is the most common language in South Africa, followed closely by the Xhosa language group (Statistics South Africa 2012). The Zulu and Xhosa people live predominantly in the Kwazulu Natal and Eastern Cape provinces respectively, in the south and south-eastern regions of southern Africa (Figure 5). Although these regions are where the highest overall WFP species richness increases are projected to occur, for the WFP species specifically used by these two language groups, range decreases are projected for 39% of species used by Zulu people and 43% of species used by Xhosa people under RCP 2.6, and 62% (Zulu) and 71% (Xhosa) under RCP 8.5 (Supplementary Table 5). These changes indicate the potential for substantial turnover in WFP species available for communities in these regions, and the need for local food systems and livelihoods to adapt to both losses of familiar species and gains of unfamiliar species.

For the majority of other language groups residing in southern Africa, varying degrees of WFP change are projected, with increases in species richness projected under RCP 2.6 and decreases under RCP 8.5 (Supplementary Table 5). However, WFP species used by the Southern Sotho language group, which resides mostly in Lesotho and the Free State province of South Africa, are at risk even under RCP 2.6, indicating that traditional practices related to WFPs are especially at risk for this group. In the north-western parts of southern Africa, the Khoesaan people, which includes the Ju’hoan, Khoekoe, Kxoe and Xóõ language groups, can be found in isolated locations around the dry Kalahari region of Namibia, Botswana, and South Africa (Welcome and Van Wyk 2019; 2020). This region has relatively few WFPs and the Khoesaan people mostly use WFP roots (Welcome and Van Wyk 2019; 2020). An increase in WFP species richness along the west coast of southern Africa may be beneficial to these groups, as the region is unsuitable for crop production now and into the future (Figure 6). However, WFPs used by two language groups within the Khoesaan are projected to experience considerable range decreases under RCP 8.5. These are the Xóõ and Khoekhoe, with 74% and 69% of species projected to experience range decreases (Supplementary Table 5). The Xóõ is also the only group where more than 50% of species are expected to experience range decreases under RCP 2.6. Considering that already water-stressed countries like Botswana and Namibia are likely to get even hotter, drier and more water-stressed (Hoegh-Guldberg et al. 2018), people from these two language groups should be prioritised for climate change adaptation strategies.

The projected losses in WFP diversity, especially under RCP 8.5, emphasize the urgency of documenting traditional knowledge on WFP use. The loss of traditional ecological knowledge in sub-Saharan Africa, is one of the greatest challenges among indigenous rural communities for accomplishing sustainable natural resource management and retaining cultural continuity (Rim-Rukeh et al. 2013; Bruyere et al. 2016; Ketlhoilwe and Jeremiah 2016; Maroyi and Cheikhyoussef 2017). In southern Africa, knowledge of indigenous plant use requires urgent documenting to not be lost irretrievably to future generations (Van Wyk and Gericke 2018). Furthermore, if we take into consideration the projected shifts in climatically-determined ranges for WFP species, language groups that have not traditionally used particular species may now have the opportunity to do so if WFP species are able to track shifting climate conditions. For other language groups, valuable species may be lost. It is thus essential that traditional knowledge on the use of WFPs is documented in a way that is respectful of the lifeways of local communities and in a form that enables the sharing of this knowledge beyond the current area of occurrence of a particular WFP species. This also have the potential to benefit the development of climate change adaptation strategies that are sustainable and participatory (Robinson and Herbert 2001).

The impact of climate change on agricultural crops will threaten food security by affecting the affordability and availability of nutritious food, especially in developing countries (Lloyd et al. 2011). Across sub-Saharan Africa, rates of undernutrition are already high, with global warming of 1.2–1.7°C projected to increase the undernourished proportion of the population by 25–90% by 2050 (Lloyd et al. 2011). WFPs are already an essential livelihood safety net when other sources of food fail, such as during times of drought or other natural disasters (Vinceti et al. 2013; Wunder et al. 2014; Shumsky et al. 2015) and might form part of climate change adaptation strategies for food security. However, for large regions of South Africa’s North West and Free State provinces, as well as parts of northern Namibia, both crop yield losses and species richness decreases of those WFPs most important for food security and as nutritional supplements are projected under RCP 2.6 and RCP 8.5 (Figure 6). Under RCP 8.5, the highest risk to both crops and WFPs are on the north-eastern border of South Africa and Botswana (Figures 6B and D), as well as for sorghum and WFPs in northern Namibia and Eswatini (Figure 6D). People in these regions will be less able to rely on WFPs as a nutritional safety net when crops fail under future climate conditions.

In contrast, for other regions in southern Africa, limiting global warming to below 2 °C may enable WFPs to continue to form part of a critical nutritional buffer against climate change for vulnerable communities. For example, in parts of the Eastern Cape province of South Africa, maize and sorghum yield are projected to decrease, and in northern regions of Namibia, sorghum yield losses are projected (Figure 6). In these same regions, climatic conditions are projected to become suitable for a greater number of WFP species (Figure 6). A greater diversity of WFPs may thus be able to continue to supplement the diets of rural people or farm households and provide them with essential micronutrients. However, this should not be a reason for policymakers to continue with business-as-usual, but rather that WFPs should be considered as part of an inclusive, equitable and participatory approach to climate change adaptation. If we follow an RCP 8.5 global warming pathway, WFPs are more at risk and may not be able to help people adapt to climate change.

Given the projected persistence of multiple WFPs under moderate or near-term global warming levels, encouraging the cultivation of WFPs could be a useful strategy to enhance food security, adapt to and mitigate climate change, and provide cash income for rural communities (Legwaila et al. 2011). A variety of native wild fruit tree species have already been domesticated in several eastern and southern Africa countries (Taylor et al. 1996; Akinnifesi et al. 2004, 2006, 2008; Leakey 2005; Kalaba et al. 2009) along with some medicinal plants (Mander et al. 1996). In these countries, it is also customary for farmers to leave native wild fruit trees when they clear land for growing crops (Legwaila et al. 2011). In Botswana, wild fruit tree planting in backyard gardens is quite common in both rural and urban areas (Taylor et al. 1996). In contrast, cultivation of wild vegetables has not received as much attention. Cultivation could be limited by their availability in the wild, but also due to a lack of knowledge in propagation techniques and husbandry skills (Legwaila et al. 2011). Furthermore, the monetary value of WFP harvests is also not well known (High and Shackleton 2000). In a case study of a rural village in the Mpumalanga province of South Africa, it was found that each household harvested four to five WFP species, with WFPs representing 31% of total dietary intake, and the mean monetary value of WFPs per household approximately 50% of that of cultivated crops (High and Shackleton 2000). However, this estimate also includes naturalised exotic species, highlighting their importance as food plants — especially as vegetables (Mogale et al. 2019) — and the need for their inclusion, along with indigenous species, in future research on climate change risk to WFPs. Thus, both native and exotic WFPs can be a significant source of income for rural people in southern Africa and may be even more so if cultivated. Apart from providing food and cash income, WFPs, especially wild fruit trees, have the added benefit of capturing and storing atmospheric carbon. Some WFPs have also been found to contain more nutrients than their cultivated counterparts (Smith et al. 1996; Kobori and Rodriguez Amaya 2008) and could therefore be important genetic resources for developing future crops (Jarvis et al. 2008).

## Conclusion

Knowing the potential impact of climate change on WFPs could afford valuable time for people to adapt to changing food security circumstances, especially for communities that are also vulnerable due to crop yield losses and already use WFPs as an important source of nutrition. Although we project that many WFP species are at risk of range reduction in southern Africa, even under a low warming scenario, WFPs could also play an important role in times of low agricultural yield as a result of changing climate conditions. This is especially true in low-income, rural communities that are reliant on smallholder farming, and for global warming below 2 °C above pre-industrial levels. Furthermore, the cultivation of WFPs could prove valuable for climate change adaptation planning, especially in the most vulnerable regions of southern Africa. However, in order to ensure future food security, more research is needed on WFP uses, nutritional value, responses to climate change, suitability for cultivation, and the climate change risk to naturalised exotic WFP species. By looking beyond the farm level and conventional crops to the exceptional diversity of WFPs people obtain from their environment, this research is a step towards the goal of understanding the linkages between WFPs, traditional knowledge, food security, climate change, and agriculture.

## Acknowledgements

C.W. and C.H.T. acknowledge support from the FLAIR Fellowship Programme: a partnership between the African Academy of Sciences and the Royal Society funded by the UK Government’s Global Challenges Research Fund. C.M. acknowledges funding from National Science Foundation grant DBI-1913673 and HDR-1934712. We acknowledge B-E van Wyk and AK Welcome for supplying the food plant database in the form of an excel spreadsheet. We also acknowledge the ISIMIP coordination team for their roles in producing, coordinating, and making available the ISIMIP model output and the modelling groups that provided the agricultural data. Thank you to M.W. Wessels for the design of the language group distribution map.

## Notes

### Competing Interest Statement

The authors have declared no competing interest.

