## Supplementary info for "Climate change risk to southern African wild food plants"

**Journal:** Regional Environmental Change

**Authors:** Carina Wessels[1]*; Cory Merow[2]; Christopher H. Trisos [1,3] **

**Affiliations:** [1] African Climate and Development Initiative, University of Cape Town, South Africa. [2] Ecology and Evolutionary Biology, University of Connecticut, Storrs, CT 06268, USA.[3] Centre for Statistics in Ecology, the Environment and Conservation, University of Cape Town, South Africa.

**Running title:** Climate change risk to southern African wild food plants

Supplementary Table 1. List of categories of wild food plant (WFP) use, and known indigenous language/ethnic groups that make use of WFPs, in southern Africa (Welcome and Van Wyk 2019, 2020).

| Categories of WFP use | Language/ethnic groups |
|---|---|
| Alcoholic beverage | Herero |
| Coffee substitute | Ju'hoan |
| Cooked vegetables | Kalanga |
| Cooking & other oils | Khoekhoe |
| Famine food | Kwangali |
| Flavourant | Kxoe |
| Ground as flour | Lozi |
| Milk curdler | Manyo |
| Moisture/thirst quencher | Mbukushu |
| Non-alcoholic beverage | Northern Sotho |
| Preservative | Southern Sotho |
| Savoury preserve | Swati |
| Snack (fresh/in situ) | Tsonga |
| Suppressant | Tswana |
| Sweet preserve | Venda |
| Syrup | Wambo |

| | |
|---|---|
| Tea substitute | Xhosa |
| Unknown | Xóõ |
| Yeast | Zulu |

Supplementary Table 2. The seven Global Climate Models (and their abbreviations) from which data on future climate projections were used for this study.

| Global Climate Model | Abbreviation | Reference |
|---|---|---|
| Beijing Climate Center, Climate System Model, version 1.1 | BCC-CSM1.1 | Xin et al. 2013 |
| Community Climate System Model, version 4 | CCSM4 | Gent et al. 2011 |
| Centre National de Recherches Météorologiques Coupled Global Climate Model, version 5 | CNRM-CM5 | Voldoire et al. 2013 |
| Geophysical Fluid Dynamics Laboratory Climate Model, version 3 | GFDL CM3 | Griffies et al. 2011 |
| Hadley Centre Global Environment Model, version 2, Earth System | HadGEM2-ES | Collins et al. 2011 |
| Model for Interdisciplinary Research on Climate, Earth System Model | MIROC-ESM | Watanabe et al. 2011 |
| Max Planck Institute Earth System Model, low resolution | MPI-ESM-LR | Giorgetta et al. 2013 |

Supplementary Table 3. The four crop models and five Global Climate Models used to project future crop yield change under RCP 2.6 and RCP 8.5. All models were used in projections for maize and sorghum, except GEPIC and LPJ-GUESS that were not available for sorghum.

| Crop model | Abbreviation | Reference |
|---|---|---|
| Environmental Policy Integrated Climate model, run by the University of Natural Resources and Life Sciences Vienna (BOKU) | EPIC-BOKU | Kiniry et al. 1995 |
| Geographic Information System-based Environmental Policy Integrated Climate Model | GEPIC | Folberth et al. 2012 |

| | | |
|---|---|---|
| Integrated Model to Assess the Global Environment | IMAGE | Bouwman et al. 2006 |
| Lund Potsdam Jena General Ecosystem Simulator | LPJ-GUESS | Lindeskog et al. 2013 |
| **Global Climate Model** | **Abbreviation** | **Reference** |
| Geophysical Fluid Dynamics Laboratory Earth System Model with MOM, version 4 component | GFDL-ESM2M | Dunne et al. 2012; 2013 |
| Hadley Centre Global Environment Model, version 2, Earth System | HadGEM2-ES | Collins et al. 2011 |
| L'Institut Pierre-Simon Laplace Coupled Model, version 5A, low resolution | IPSL-CM5A-LR | Dufresne et al. 2013 |
| Model for Interdisciplinary Research on Climate, Earth System Model, Chemistry Coupled | MIROC-ESM-CHEM | Watanabe et al. 2011 |
| Norwegian Earth System Model, version 1 (intermediate resolution) | NORESM1-M | Bentsen et al. 2013 |

Supplementary Table 4. The percentages of wild food plant species, for each of 19 categories of plant use, that are projected to either experience a range loss or gain under RCP 2.6 and RCP 8.5. The numbers of species (n) are shown for each category of plant use. Where the total percentage of species with range loss or range gain for a given scenario is greater than 50%, this is shown in bold.

| | **% of species with range loss** | | | | **% off species with range gain** | | | |
|---|---|---|---|---|---|---|---|---|
| | **RCP 2.6 range loss (%)** | | **RCP 8.5 range loss (%)** | | **RCP 2.6 range gain (%)** | | **RCP 8.5 range gain (%)** | |
| | **0-25%** | **>25%** | **0-25%** | **>25%** | **0-25%** | **>25%** | **0-25%** | **>25%** |
| All species (n=1190) | 37 | 3 | **44** | **22** | **40** | **20** | 18 | 16 |
| Categories of use important during famine: | | | | | | | | |
| Famine food (n=64) | 39 | 2 | **45** | **14** | **38** | **22** | 22 | 19 |
| Ground as flour (n=91) | 37 | 5 | **40** | **27** | **40** | **18** | 18 | 15 |
| Cooked vegetables (n=407) | 39 | 4 | **44** | **24** | **40** | **18** | 17 | 14 |
| Snack (fresh/in situ) (n=686) | 37 | 3 | **41** | **22** | **38** | **23** | 19 | 18 |
| Other categories of use: | | | | | | | | |
| Alcoholic beverage (n=56) | 30 | 2 | 30 | 14 | **36** | **32** | **34** | **21** |
| Coffee substitute (n=22) | 44 | 5 | **27** | **36** | **33** | **18** | 18 | 18 |
| Cooking & other oils (n=13) | 38 | 0 | **23** | **31** | **31** | **31** | 23 | 23 |
| Flavourant (n=86) | 35 | 2 | **43** | **22** | **40** | **23** | 19 | 16 |
| Milk curdler (n=35) | **51** | **6** | **46** | **29** | 26 | 17 | 11 | 14 |

| | | | | | | | | |
|---|---|---|---|---|---|---|---|---|
| Moisture/thirst quencher (n=61) | 34 | 0 | **46** | **31** | **49** | **16** | 16 | 7 |
| Non-alcoholic beverage (n=49) | 33 | 6 | **22** | **29** | **24** | **37** | 22 | 27 |
| Preservative (n=30) | 49 | 0 | **47** | **27** | **38** | **13** | 20 | 7 |
| Savoury preserve (n=19) | 37 | 0 | **42** | **21** | **42** | **21** | 16 | 21 |
| Suppressant (n=11) | 36 | 9 | **55** | **36** | **45** | **9** | 0 | 9 |
| Sweet preserve (n=49) | 37 | 0 | **35** | **24** | **35** | **29** | 22 | 18 |
| Syrup (n=6) | **67** | **0** | **50** | **33** | 17 | 17 | 0 | 17 |
| Tea substitute(n=71) | 45 | 0 | **54** | **27** | **44** | **11** | 14 | 6 |
| Unknown (n=164) | 38 | 1 | **47** | **18** | **45** | **16** | 21 | 14 |
| Yeast (n=36) | **53** | **0** | **42** | **36** | 33 | 14 | 11 | 11 |

Supplementary Table 5. The percentages of wild food plant species used by each of 19 language groups that are projected to either experience a range loss or a range gain, under RCP 2.6 and RCP 8.5. The numbers of species (n) are shown for each language group. Where the total percentage of species with range loss or range gain for a given scenario is greater than 50%, this is shown in bold.

| | **% of species with range loss** | | | | **% off species with range gain** | | | |
|---|---|---|---|---|---|---|---|---|
| | **RCP 2.6 range loss (%)** | | **RCP 8.5 range loss (%)** | | **RCP 2.6 range gain (%)** | | **RCP 8.5 range gain (%)** | |
| | 0-25% | >25% | 0-25% | >25% | 0-25% | >25% | 0-25% | >25% |
| Herero (n=377) | 34 | 5 | **34** | **26** | **34** | **27** | 19 | 20 |
| Ju'hoan (n=287) | 28 | 6 | **30** | **25** | **37** | **29** | 22 | 23 |

| | | | | | | | | |
|---|---|---|---|---|---|---|---|---|
| Kalanga (n=288) | 31 | 5 | **36** | **23** | **40** | **25** | 20 | 20 |
| Khoekhoe (n=452) | 39 | 4 | **41** | **28** | **36** | **21** | 16 | 15 |
| Kwangali (n=94) | 28 | 6 | **29** | **30** | **35** | **31** | 23 | 18 |
| Kxoe (n=250) | 26 | 4 | **31** | **20** | **37** | **32** | 23 | 26 |
| Lozi (n=181) | 23 | 6 | 27 | 20 | **35** | **36** | **24** | **29** |
| Manyo (n=112) | 24 | 6 | **28** | **26** | **36** | **34** | 21 | 26 |
| Mbukushu (n=198) | 28 | 5 | **28** | **25** | **36** | **31** | 22 | 25 |
| NSotho (n=747) | 38 | 2 | **47** | **16** | **39** | **20** | 21 | 16 |
| SSotho (n=615) | 47 | 2 | **53** | **23** | **42** | **10** | 18 | 6 |
| Swati (n=748) | 37 | 1 | **48** | **13** | **41** | **20** | 22 | 16 |
| Tsonga (n=787) | 37 | 1 | **46** | **13** | **39** | **23** | 22 | 19 |
| Tswana (n=790) | 38 | 3 | **43** | **21** | **38** | **21** | 20 | 15 |
| Venda (n=622) | 36 | 2 | **44** | **13** | **38** | **25** | 23 | 20 |
| Wambo (n=222) | 32 | 7 | **32** | **27** | **34** | **28** | 21 | 20 |
| Xhosa (n=695) | 42 | 0 | **53** | **18** | **44** | **14** | 19 | 10 |
| Xóõ (n=106) | **42** | **8** | **28** | **45** | 30 | 19 | 13 | 13 |
| Zulu (n=856) | 38 | 1 | **46** | **16** | **40** | **20** | 22 | 16 |
